# *C. elegans* aversive olfactory learning generates diverse intergenerational effects

**DOI:** 10.1101/2020.02.07.939017

**Authors:** Ana Gonçalves Pereira, Xicotencatl Gracida, Konstantinos Kagias, Yun Zhang

## Abstract

Parental experience can modulate the behavior of the progeny through the inheritance of phenotypic traits acquired by the progenitors. While the molecular mechanisms for behavioral inheritance are studied under several environmental conditions, it remains largely unexplored how the nature of the parental experience affects the information transferred to the next generation. To address this question we used *C. elegans*, a nematode that feeds on bacteria in its habitat. Some of these bacteria are pathogenic and the worm learns to avoid them after a brief exposure. We found, unexpectedly, that a short parental experience increased the preference for the pathogen in the progeny. Furthermore, increasing the time of parental exposure switched the response of the progeny from attraction to avoidance. To characterize the underlying molecular mechanisms, we found that the RNA-dependent RNA Polymerase (RdRP) RRF-3, required for the biogenesis of 26G endo-siRNAs, regulated both types of intergenerational effects. Together, we show that different parental experiences with the same environmental stimulus generate diverse effects on the behavior of the progeny through small RNA-mediated regulation of gene expression.

## INTRODUCTION

In many organisms, the parental experience modulates the behavior and physiology of the progeny in such a way that the progeny phenocopy the acquired phenotype of the parents (Agrawal, Laforsch, and Tollrian 1999; Remy 2010; Rechavi et al. 2014; Bozler, Kacsoh, and Bosco 2019; Dias and Ressler 2014; Gapp et al. 2014; Ng et al. 2010). This form of plasticity “anticipates” that the next generation will experience a similar environment to their parents, in which case it may prove to be adaptive. However, very often the progeny is exposed to environmental conditions distinct from their progenitors and under these circumstances the same phenotypic traits that were adaptive to the parents become maladaptive to the progeny (Painter, Roseboom, and Bleker 2005; Wells 2007). In addition, the same parental experience can affect the progeny differently depending on their sex, genotype, further ancestral history and life experiences (Deas, Blondel, and Extavour 2019; Palominos et al. 2017; Crews et al. 2014; Dew-Budd, Jarnigan, and Reed 2016; Bohacek and Mansuy 2015). These results suggest that inter- and transgenerational regulations are influenced by many different factors and do not always phenocopy the parental acquired traits.

One of the factors that remain largely unexplored is how the duration, intensity or frequency of the parental experience affects the modulation of certain traits in the progeny (Öst et al. 2014; Palominos et al. 2017; Kundakovic et al. 2013; Mueller and Bale 2006). Here, we examine this question using *C. elegans*, a nematode that feeds on different species of bacteria, such as the *Pseudomonas* genus that are abundant in the natural habit of the nematode (Samuel et al. 2016). However, certain *Pseudomonas* strains, such as *Pseudomonas aeruginosa* PA14, infect the worm after being ingested which results in a slow death over the course of days (Tan, Mahajan-Miklos, and Ausubel 1999). Therefore, *Pseudomonas aeruginosa* PA14 is food and pathogen, both of which are critical for survival.

We have previously shown that adult *C. elegans* robustly learns to reduce their preference for the odorants of PA14 after transiently feeding on the pathogen for 4 hours (Zhang, Lu, and Bargmann 2005; Ha et al. 2010a). This form of learning is contingent on the pathogenesis of the training bacteria and resembles the Garcia effect, a robust form of conditioned aversion that allows animals to learn to avoid the smell or taste of a food that makes them ill (Garcia, Hankins, and Rusiniak 1974; Zhang, Lu, and Bargmann 2005; Ha et al. 2010a).

Because ingesting the pathogenic food PA14 can be lethal to the worm, the intergenerational transfer of information regarding the valence of cues that signal this food source could be adaptive for the next generation. Therefore, we asked whether learning to avoid PA14 in the adult *C. elegans* regulates the olfactory response in the progeny. Furthermore, we asked whether the parental learning experiences of different duration differ in their effects on the behavioral response of the progeny.

To elucidate the molecular pathways underlying behavioral changes in progeny triggered by parental experience, we characterized the small non-coding RNA pathways that were implicated in inter- and trans-generational inheritance in several species (Posner et al. 2019; Rechavi et al. 2014; Moore et al. 2019; Palominos et al. 2017; Brennecke 2008; Gapp et al. 2014; Bohacek and Mansuy 2015; Okamura and Lai 2008; Bond and Baulcombe 2014). These pathways haven been extensively studied in *C elegans* and three major classes of small RNAs have been described depending on their size, function and biogenesis: small interfering RNAs (endo-siRNAs and exo-siRNAs for endogenously and exogenously produced small RNAs, respectively), microRNAs, and PIWI-interacting RNAs (piRNAs). Endo-siRNAs are encoded in the worm genome and target the host transcripts; the RNA-dependent RNA Polymerase (RdRP) RRF-3 uses the endogenous mRNAs as templates to synthesize the complementary strand. The resulting double-strand RNAs (dsRNAs) are further cleaved into 26G endo-siRNAs that target genes expressed in both the soma and the germline (Gent et al. 2009; Han et al. 2009; Vasale et al. 2010; Rankin 2015; Claycomb et al. 2009). Mutating RRF-3 leads to downregulation of the 26G endo-siRNAs in the sperm, oocytes and embryos and upregulation of the expression of many genes (Han et al. 2009; Gent et al. 2010; Lee, Hammell, and Ambros 2006). The importance of this pathway in epigenetic inheritance in response to environmental stimuli has been recently reported in a study documenting the transgenerational inheritance of endo-siRNAs in response to starvation (Rechavi et al. 2014).

The piRNA pathway has also been implicated in several aspects of epigenetic inheritance in response to the environmental stress (Belicard, Jareosettasin, and Sarkies 2018; Moore et al. 2019; Gapp et al. 2014). piRNAs are 21 to 22-nucleotide RNAs that are enriched in the germline where they protect the integrity of the genome and regulate the development of the germ cells. The piRNAs form complexes with the PIWI-clade Argonauts, such as PRG-1 and PRG-2, to target both the endogenous and the exogenous transcripts (Das et al. 2008; Ashe et al. 2012; Youngman and Claycomb 2014).

Given the documented role of these two pathways in epigenetic inheritance in the context of an environmental challenge, we characterized the function of RRF-3 and PRG-2 in the aversive learning of PA14 in the adult animals, and the effects of the learning experience on the olfactory response of the progeny to PA14. First, we found that training adult *C. elegans* with PA14 for 4 hours reduced the preference for the odorants of the pathogen and increased the preference for PA14 in the progeny. Mutating either *rrf-3* or *prg-2* abolished the parental training-induced change in the olfactory preference of the progeny, without compromising the learning ability of the mothers. Increasing the training duration from 4 hours to 8 hours, while generating similar learning in the mothers, switches the preference of the progeny from attractive to aversive and the parental experience-induced behavioral change also depends on *rrf-3*. Together, our results show that parental learning experiences of different duration generate different effects in the behavior of the progeny and endo-siRNA pathways regulate these intergenerational effects.

## MATERIALS AND METHODS

### Experiment model and subject details

*C. elegans* hermaphrodites were used in this study and cultivated using the standard conditions (Brenner 1974). The strains used in this study include: N2 (Bristol), ZC2834 *rrf-3(pk1426)* II, YY13 *rrf-3(mg373)* II, ZC2987 *prg-2(n4358)* IV, WM162 *prg-2(tm1094)* IV, CX4998 kyIs140 I; *nsy-1(k397)* II.

### Bacterial Strains

For this study we used the non-pathogenic bacteria *Escherichia coli* OP50 (*Caenorhabditis* Genetics Center (CGC)) and the pathogenic bacteria *Pseudomonas aeruginosa* PA14 (Kim et al. 2002).

### Method details

#### The aversive olfactory training of P0s with PA14 and the cultivation of F1s

Training of P0s with PA14 and the learning test were preformed mainly as previously described (Ha et al. 2010a; Zhang, Lu, and Bargmann 2005) with minor modifications. To prepare the training plates, individual OP50 colonies or PA14 colonies were used to inoculate 50 mL nematode growth medium (NGM, 2.5g/L Bacto Peptone, 3.0g/L NaCl, 1mM CaCl2, 1mM MgSO4, 25mM KPO4 pH6.0, 5mg/L cholesterol), which were cultivated with shaking at 27°c overnight. 800 μL (for 4-hour training protocol) or 400 μL (for 8-hour training protocol) OP50 or PA14 culture was spread onto each 10 cm NGM plate, which was incubated at 27°C for 2 days to prepare the naive control and the training plates, respectively. To prepare the plates where P0s, F1s and F2s were cultivated (standard conditions), individual colonies of OP50 were inoculated in 50 mL of Luria-Bertani (LB) medium and cultivated with shaking at 27°C overnight. To make a bacterial lawn, 400 μL OP50 culture was spread in 10 cm NGM plate and incubated for at least 24 hours under the room temperature. To perform training and control, eggs were extracted from first-day adult hermaphrodites cultivated under the standard conditions on the benign bacteria *Escherichia coli* OP50 (Brenner 1974) and grown until they reached adulthood. Populations of synchronized young adults were transferred to the training plate or the control plate and kept at 20°C for 4 or 8 hours as specifically described. By the end of the training, a group of naive control and trained P0 worms were randomly picked to measure the olfactory choice and learning in the automated assay. The rest of the P0 worms were collected from the plates with S-basal medium and passed through a cell strainer 40-micrometer nylon filter (Falcon) to remove laid eggs. The collected P0 worms were then treated with a bleach solution to isolate F1 embryos, which were hatched and cultivated under the standard conditions with OP50 as food until the adult stage.

#### Olfactory preference assay using the automated olfactory assay

The automated assay (Droplet assay) that quantified the olfactory preference in individual animals was performed as previously described (Ha et al. 2010a) with some modifications. Briefly, by the end of the training, several naive and trained P0s were randomly picked and washed with buffer and individually placed in the droplets of 2 μL NMG buffer (1mM CaCl2, 1mM MgSO4, 25mM KPO4 pH6.0) in an enclosed chamber and subjected to alternating airstreams that were odorized with OP50 or PA14 by blowing clean air through the supernatant of freshly generated bacterial cultures. Each olfactory stimulation lasted for 30 seconds and each assay contained 12 cycles of stimulation. The locomotion of the worms was video recorded and the large body bends of the tested worms were identified with machine-vision softwares. Because large body bends are followed by reorientation, a higher rate of large body bends indicates a lower preference for the tested airstream (Pierce-Shimomura, Morse, and Lockery 1999; Ha et al. 2010b). The choice index (CI) of each worm was defined as the body bends evoked by OP50 smell minus the body bends evoked by PA14 smell and normalized by the total body bends [CI = (bends evoked by OP50 − bends evoked by PA14)/(bends evoked by OP50 + bends evoked by PA14)]. Each assay examined 5 – 6 naive worms and 5 – 6 trained worms in parallel. The learning index (LI) was defined as the median value of CI of the naive worms minus the median value of CI of the trained worms (LI = CI of naive worms – CI of trained worms). A positive LI indicates learned avoidance of PA14.

#### Olfactory preference assay using the two-choice assay

The two-choice plate assay is similar to the one described in Zhang et al., 2005, except for several modifications. To measure the olfactory preference for bacteria, a drop of 5 μL supernatant of OP50 culture in NGM medium and a drop of 5 μL PA14 culture in NGM medium were put 2 cm apart on a 6 cm NMG plate. In each assay, one worm was placed on the plate equidistant to the two drops of the bacteria culture supernatant right after the drops were put on the plate and allowed to crawl to the preferred stimulus. The movement and the choice of each worm were recorded and later analyzed. The worms that did not make choice after 10 minutes were also counted in the total number. The worms that disappeared during the assay (because they moved to the side of the plate) were not included in the total number. The parental experience-induced choice index in the two-choice assay was defined as the number of worms that chose PA14 minus the number of worms that chose OP50 normalized by the total number of worms tested [parental experience-induced choice index = (number of worms that choose PA14 - number of worms that choose OP50)/total number of worms] and the parental experience-induced learning index for the two-choice assay was defined as the parental experience-induced choice index of F1s from naive P0s (F1 WT OP50_mothers_) minus parental experience-induced choice index of F1s from trained P0s (F1 WT PA14_mothers_). A positive parental experience-induced learning index indicates increased avoidance of PA14 in F1s induced by the parental experience with PA14.

#### Slow killing assay

The slow killing assay was performed essentially as previously described (Tan, Mahajan-Miklos, and Ausubel 1999). 200 μL freshly prepared PA14 culture incubated at 27°C in the LB medium overnight were spread into a 4 cm diameter lawn on a 6 cm NMG plate and incubated at 37°C for 24 hours and then left at room temperature for another 24 hours before the assay. 20 F1 young adult hermaphrodites were transferred onto each slow killing plate and kept at 25°C. The dead and the alive worms were counted at the specific time points as shown in the figure.

#### The analysis of parental effects induced by 8-hour training

To separate the experiments in which P0s exhibited high learning indexes from the experiments in which P0s exhibited low learning indexes after 8-hour training with PA14, we plotted the values of learning index of P0s against the values of Parental experience_induced learning index of the respective progeny and tested for a significant linear correlation between the variables using a Permutation test where labels were exchanged for 5,000 times. We then calculated the average P0 learning index to separate the experiments into two groups - the experiments in which the P0 learning indexes were higher than the average learning index, and the experiments in which the P0 learning indexes were smaller than the average learning index (Figure 2E).

#### Quantification of the body size and the chemotactic movements

The body size and the parameters of the chemotactic movements in the two-choice assays were quantified by analyzing the recorded worms with the Wormlab tracker (https://www.mbfbioscience.com/wormlab).

### Quantification and statistical analysis

All data analyzes were conducted using Matlab_R2015b. The statistical methods, sample size and number of the replicates are indicated in the legend of each figure.

## RESULTS

To examine the pattern through which parental experience modulates offspring behavior, we exposed *C. elegans* to the pathogenic bacteria *Pseudomonas aeruginosa* strain PA14 and asked whether maternal aversive learning with PA14 leads to a decreased attraction towards the pathogen in the adult progeny.

We cultivated the worms under the standard conditions on the benign bacteria strain *Escherichia coli* OP50 until the adult stage (Brenner 1974). We then trained a group of the young adult hermaphrodite mothers (P0s) by feeding them on PA14 for 4 hours and fed the rest of the worms with OP50 in parallel as naive controls (Figure 1A, Materials and Methods). We quantified the preference between the smell of OP50 and PA14 in naive and trained mothers using the droplet assay, in which individual worms swim in droplets of buffer to which the odorants of the tested bacteria are delivered with air streams (Ha et al. 2010b). The embryos from the rest of the mothers were harvested with a bleach solution that dissolved P0 bodies and the associated bacteria, thus preventing the progeny from a direct contact with PA14. The F1 embryos were cultivated on *E. coli* OP50 until they reached adulthood at which stage their preference between the smell of OP50 and the smell of PA14 was tested using a two-choice assay on plate. In this test, an individual worm navigates to choose between a drop of supernatant of an OP50 or PA14 culture immediately after the supernatants were placed on the plate (Figure 1A). We recorded the behavior of the F1 worms during the two-choice assay in order to quantify specific behavioral features that may underlie changes in olfactory chemotaxis (H. Liu et al. 2018). While the droplet assay allowed us to rapidly and systematically measure the olfactory preference of several P0s at the same time before handling the F1 embryos, the two-choice assay allowed us to examine the odorant-guided movements of the progeny in details (Materials and Methods). In both types of assays, a positive choice index indicates a preference for PA14, and a positive learning index indicates that training with PA14 in P0s reduces the preference for PA14 in comparison with naive controls (Figure 1A).

**Figure 1.**
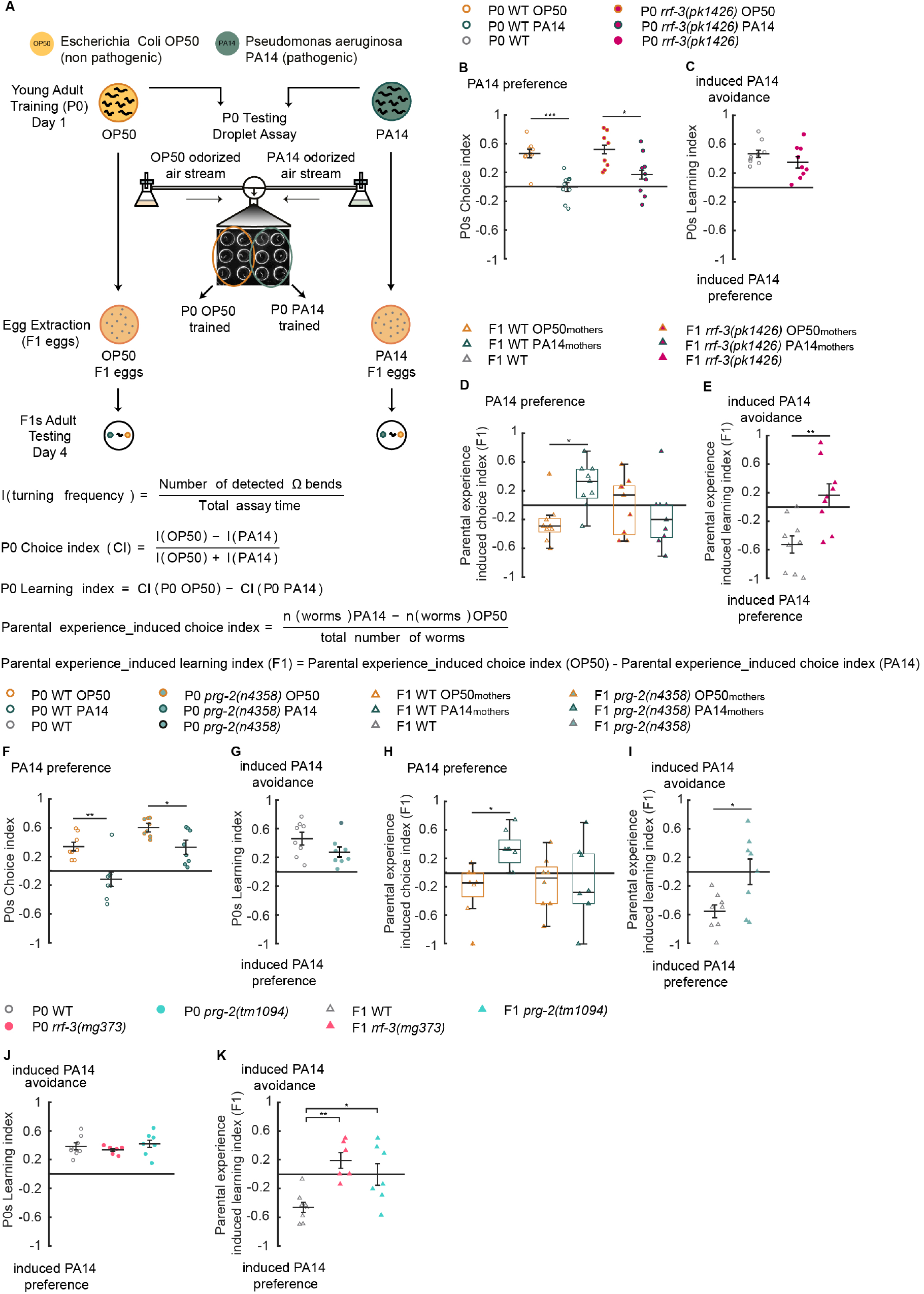
Aversive training with pathogenic bacterium PA14 for 4 hours increases offspring olfactory preference for PA14, in a small RNA pathway dependent manner. **(A)** Schematics for the training procedure and for the assays to quantify olfactory learning in P0s and parental experience-induced olfactory learning in F1s (Materials and Methods). **(B, C)** Naive wild-type (WT) animals fed on *E. coli* OP50 (P0 WT OP50) prefer the odorants of PA14 (**B**), while training with PA14 (P0 WT PA14) decreases the preference (**B, C**). The deletion mutation *rrf-3(pk1426)* does not alter PA14 aversive learning in P0 mothers (**B, C**). A two-tailed Student’s *t*-test with Bonferroni correction for multiple comparisons was used in **(B)** (n = 9 biological replicates each; *** p < 0.001, * p < 0.05). A two-tailed student’s *t*-test was used for comparisons in **(C)** (n = 9 biological replicates each, p > 0.05). **(D, E)** The progeny of the WT trained mothers (F1 WT PA14_mothers_) show an increased preference for PA14 in comparison with the progeny of the WT naive mothers (F1 WT OP50_mothers_). The deletion mutation *rrf-3(pk1426)* abolishes this increase (**D**). WT F1s generate a negative average parental experience-induced learning index, which contrasts with the parental experience-induced learning index in the F1s of *rrf-3(pk1426)* mutants (**E**). A Kruskal-Wallis test with Tukey’s post hoc comparison was used for comparisons in **(D)** (n = 9 biological replicates each; Chi-sq = 9.3; F1 WT OP50_mothers_ *vs* F1 WT PA14_mothers_, * p < 0.05; for all other comparisons, p > 0.05). A two-tailed student’s *t*-test was used for comparisons in **(E)** (n = 9 biological replicates each, ** p < 0.01). (**F, G**) The *prg-2(n4358)* mutant mothers (P0) learn to avoid the pathogen similarly to WT. A two-tailed Student’s *t*-test with Bonferroni correction for multiple comparisons was used in **(F)** (n = 8 biological replicates each; ** p < 0.01, * p < 0.05). A two-tailed student’s *t*-test was used for comparisons in **(G)** (n = 8 biological replicates each, p > 0.05). (**H, I**) The learning experience of the mothers does not alter the olfactory preference in *prg-2(n4358)* F1s (**H**), which display parental experience-induced learning indexes significantly more positive than that of WT (**I**). A Kruskal-Wallis test with Tukey’s post hoc comparison was used for comparisons in **(H)** (n = 8 biological replicates each; Chi-sq = 9.73; F1 WT OP50_mothers_ *vs* F1 WT PA14_mothers_ * p < 0.05; for all other comparisons p > 0.05). A two-tailed student’s *t*-test was used for comparisons in **(I)** (n = 8 biological replicates each, * p < 0.05). (**J, K**) The *rrf-3(mg373)* and the *prg-2(tm1094)* mutant mothers learn in a similar way as WT P0s (**J**) but show a defective parental experience-induced learning in F1s (**K**). A one way analysis of variance ANOVA with Tukey’s post hoc comparison was used for comparisons in (**J**) (n = 8, n = 6 and n = 7 biological replicates for WT, *rrf-3(mg373)* and *prg-2(tm1094)*, respectively; p > 0.05, F = 0.568) and (**K**) (n = 8, n = 6 and n = 7 biological replicates for WT, *rrf-3(mg373)* and *prg-2(tm1094)* respectively; F = 8.99; F1 WT *vs* F1 *rrf-3(mg373)* ** p < 0.01; F1 WT *vs* F1 *prg-2(tm1094)* * p < 0.05; F1 *rrf-3(mg373) vs* F1 *prg2(tm1094)* p > 0.05). Data is normally distributed in **B, C, E, F, G, I, J** and **K** and not normally distributed in **D** and **H**. In graphs **B, C, E, F, G, I, J** and **K** horizontal bar represents Mean ± SEM and in box plots (**D** and **H)** dark horizontal bar represents median, box limits represent first and third quartile, and whiskers extend to values within 2.7 standard deviations. *** p < 0.001, ** p < 0.01, * p < 0.05. All data are included in the analysis.

### Transient parental exposure to PA14 increases the preference for the pathogen in the progeny

Consistent with our previous findings (Zhang, Lu, and Bargmann 2005; Ha et al. 2010a), mothers trained for 4 hours with PA14 (P0 WT PA14) showed a reduced preference for the odorants of the pathogen in comparison to naïve mothers (P0 WT OP50). This difference is indicated by a positive P0 Learning index (Figure 1B and 1C, left side of the graphs). We also analyzed the preference of P0s using the two-choice assay on plate and found that both types of assays demonstrated similar training effects (Supplementary Figure 1A). Next, we examined whether the learning experience of the P0s modulated the olfactory behavior of their progeny. We measured the difference between the choice index of the progeny of the naive (F1 WT OP50_mothers_) and the trained (F1 WT PA14_mothers_) mothers, and defined it as the Parental experience_induced learning index (F1) (Figure 1A). In contrast with P0s, we found that the progeny of trained mothers showed an increased preference for PA14 in comparison with the progeny of the naive mothers (Figure 1D, left side of the graph). This generated a Parental experience_induced learning index (F1) with a negative average value (Figure 1E, left side of the graph), which resulted from an average preference for PA14 in the progeny of the trained mothers and an average preference for OP50 in the progeny of the naive mothers (Figure 1D). This surprising result suggests that the food choices of the progeny are modulated by parental experience and that a short exposure to the pathogen PA14 reduces the preference for the odorants of PA14 in mothers, but increases this preference in the progeny.

### The endo-siRNA and piRNA pathways regulate the modulation of behavioral responses in the progeny

Next, we sought the signaling mechanisms whereby parental experience mediates the changes in the behavioral preference of the progeny. We asked if disrupting the biogenesis of the 26G endo-siRNAs affected the modulation of the olfactory choice in F1s. To this end we tested animals with a loss of function mutation in *rrf-3(pk1426)* (Simmer et al. 2002) and found that the mutant mothers displayed the normal learning after 4-hour training with PA14 (Figure 1B and 1C, right side of the graphs). However, this learning experience did not alter the olfactory preference of their progeny, demonstrated by the similar choice indexes in *rrf-3(pk1426)* F1s from the naive and the trained mothers (Figure 1D, right side of the graph) and a Parental experience_induced learning index (F1) significantly different from that of wild type (Figure 1E, right side of the graph). To further confirm the role of *rrf-3*, we tested another independently generated mutant allele, *mg373*, that harbors a missense mutation for a conserved catalytic residue (Pavelec et al. 2009). We found that *rrf-3(mg373)* mutant animals showed a similar phenotype as the *rrf-3(pk1426)* deletion mutants in both P0s and the F1 progeny (Figure 1J and 1K; Supplementary Figure 2A, 2B, 2D and 2E). Together, these results indicate that the *rrf-3*-mediated endo-siRNA pathway regulates the modulation of offspring olfactory preference by parental experience.

We next examined the involvement of the piRNA pathway that mainly regulates gene silencing and germline maintenance (Weick and Miska 2014). The piRNA population has been shown to be altered by exposure to environmental stressors, such as *Pseudomonas aeruginosa* bacteria (Belicard, Jareosettasin, and Sarkies 2018), which suggests a potential role of piRNAs in regulating parental experience-induced behavioral changes to PA14. While the PIWI-clade Argonaut *prg-1* has been previously implicated in regulating a form of pathogen avoidance that lasts for several generations (Moore et al. 2019), very little is know about the function of its homolog *prg-2* (Kasper, Gardner, and Reinke 2014; Youngman and Claycomb 2014). We tested a deletion mutation, *n4358*, in the *prg-2* gene. At least three alternatively spliced transcripts are predicted from the genomic locus of *prg-2* and the deletion in *n4358* affects all the transcripts. Two of the predicted transcripts encode pseudogenes and the third transcript encodes the homologous protein of the *C. elegans* Argonaute/Piwi-related protein PRG-1. We detected the third transcript in a wild-type cDNA library (Supplementary Figure 2G and 2H). PRG-1 regulates 21 U-RNAs (piRNAs) to mediate gene silencing and germline integrity (Bagijn et al. 2012; Batista et al. 2008; Cox et al. 1998). The putative PRG-2 protein is nearly identical to PRG-1 and the *prg-2* transcripts are found primarily in the germline cells (Hashimshony et al. 2015). We found that *prg-2* P0s exhibited a significantly decreased preference for PA14 after training and generated learning indexes comparable to wild type (Figure 1F and 1G). However, the learning experience of the *prg-2* mothers did not modulate the preference of their progeny in the same way as in wild type. The F1s of the naive and the trained *prg-2(n4358)* mothers displayed similar choice indexes for PA14, which generated the parental experience-induced learning indexes significantly different from wild type and with an average value close to 0 (Figure 1H and 1I). Furthermore, an independently generated deletion mutation, *tm1094*, in *prg-2* similarly disrupted the Parental experience_induced learning index (F1) (Figure 1J, 1K, Supplementary Figure 2C and 2F). Taken together, these findings suggest the function of the piRNA pathway in transducing parental learning experience to the progeny and modulating offspring behavior.

### Extending the duration of parental exposure to PA14 switches attraction to aversion in the progeny

We were intrigued by the increased preference for the pathogenic bacteria PA14 in the progeny of the trained mothers. We asked whether stabilizing the parental environment by extending parental training with PA14 would generate a different offspring response. We found that training mothers with PA14 for 8 hours also induced an aversive learning of PA14 (Figure 2A and 2B). However, the preference for PA14 in the progeny of the trained mothers was similar to that in the progeny of the naive mothers, which produced Parental experience_induced learning index (F1) with a mean value close to zero (Figure 2C and 2D). One feature in the choice indexes and the learning indexes of these progeny is their widespread values. This observation prompted us to examine the correlation between the learning of the mothers and the behavior of their progeny. We plotted the Parental experience-induced learning indexes in F1s (y-axis) as a function of the learning indexes of their mothers in each independent experiment and found a significant correlation. The more the mothers learned to avoid PA14, the more their offspring avoided the pathogen (Figure 2E). We calculated the average learning index of the P0s and divided all the experiments in 2 groups - “P0 low learning index” and “P0 high learning index” (Figure 2E, Material and Methods). This analysis showed that the F1s in the experiments where mothers learned to avoid PA14 at a lower level, *i.e.* “P0 low learning index”, have a negative average value for Parental experience_induced learning index (F1) (Figure 2E). This indicates that in these experiments there is a tendency for the progeny from trained mothers to prefer PA14 more than the progeny from naive mothers, similarly to the progeny from the 4-hour trained mothers. In contrast, the F1s in the experiments where mothers learned to strongly avoid PA14, *i.e.* “P0 high learning index”, have a positive average value for Parental experience_induced learning index (F1) (Figure 2E). This value indicates that in these experiments the progeny from trained mother avoids PA14 more than the progeny of naive mothers. Consistently, the F1s of the trained mothers in the “P0 high learning index” group avoided the pathogen significantly more than the F1s of the trained mothers in the “P0 low learning index” group (Supplementary Figure 3A). Together, these results show that the wide-spread behavioral response to PA14 in the progeny of the 8-hour trained mothers is modulated by differences in their mothers’ learning.

**Figure 2.**
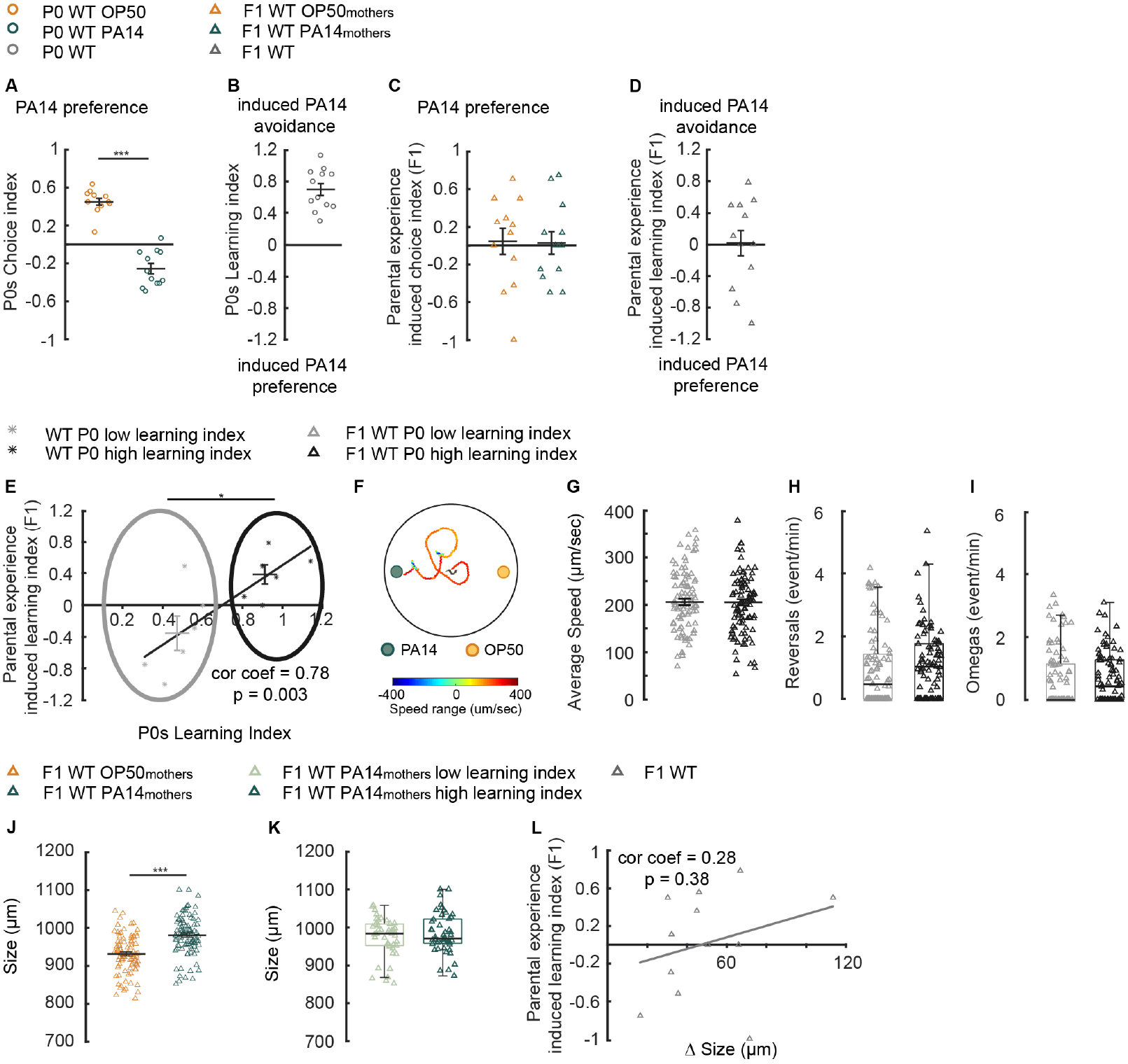
Extending parental training to 8 hours switches offspring response from attraction to aversion. **(A, B)** Training with PA14 for 8 hours decreases the preference for PA14 odorants in P0 wild-type (WT) mothers **(A)**, resulting in learned avoidance of PA14 **(B)**. A two-tailed student’s *t*-test was used for comparisons in **(A)** (n = 12 biological replicates each, *** p < 0.001). **(C, D)** The progeny of naive (F1 WT OP50_mothers_) and trained (F1 WT PA14_mothers_) mothers show similar preference for the pathogen. A two-tailed student’s *t*-test was used for comparisons in **(C)** (n = 12 biological replicates each, p > 0.05). **(E)** The parental experience-induced learning indexes in F1s are significantly correlated with the learning indexes in their mothers after 8-hour training. The progeny of the mothers in the “P0 high learning index group” (dark grey stars) display an increased avoidance of PA14; this response is distinct from the progeny of the mothers in the “P0 low learning index group” (light grey stars). A permutation test was used to test linear correlation (Materials and Methods) and a two-tailed student’s *t*-test was used for comparisons between the parental experience-induced learning indexes of the F1s from the two groups (n = 12 biological replicates, * p < 0.05). **(F)** Schematics showing the trajectory of a F1 worm in the two-choice plate assay. **(G, H, I)** The movement speed **(G)**, rate of reversals **(H)** and the rate of omega bends **(I)** for F1s in the “P0 high learning index” group and “P0 low learning index” group. A two-tailed student’s *t*-test was used for comparisons in **(G)** (p > 0.05) and a Wilcoxon rank sum test was used for comparisons in (**H, I**) (p > 0.05). n = 90 worms for F1 WT P0 low learning index and n = 93 worms for F1 WT P0 high learning index. **(J)** The effect of parental learning on the body size of the progeny. A two-tailed student’s *t*-test was used for comparisons (*** p < 0.001). n = 90 worms for F1 WT OP50_mothers_ and n = 93 worms for F1 WT PA14_mothers_. **(K)** The progeny of trained mothers have similar body size. Wilcoxon rank sum test was used for comparisons (F1 WT PA14_mothers_ low learning index *vs* F1 WT PA14_mothers_ high learning index p > 0.05; n = 45 for F1 WT PA14_mothers_ low learning index and n = 48 for F1 WT PA14_mothers_ high learning index). **(L)** The parental experience-induced learning index is not correlated with the increased size of progeny of trained mothers in comparison with progeny of naive mothers. A permutation test was used to test for linear correlation (STAR Methods). (n = 12 biological replicates). Data is normally distributed in **A - E, G** and **J** and not normally distributed in **H**, **I** and **K**. In graphs **A - E, G** and **J** horizontal bar represents Mean ± SEM and in box plots (**H**, **I** and **K)** dark horizontal bar represents median, box limits represent first and third quartile, and whiskers extend to values within 2.7 standard deviations. *** p < 0.001, * p < 0.05. All data are included in the analysis.

We then asked how the progeny from the “P0 low learning index” group and the progeny from the “P0 high learning index” group differed in their chemotactic movements in the two-choice assay. We video tracked the worms and quantified several behavioral parameters in order to address if any specific parameters were modulated by parental experience and may underlie the different food odorant choices (Figure 2F and Material and Methods). We found no difference between the F1s in the “P0 low learning index” group and the F1s in the “P0 high learning index” group in the locomotory speed, the reversal rate and the frequency of omega bends (*i.e.* the big body bends that resemble the letter Omega) (Figure 2G – 2I). These results suggest that parental learning experience with PA14 regulates the choice of the offspring without affecting these specific locomotory parameters and likely modulates offspring behavior by regulating a higher level of sensorimotor functions.

Meanwhile, we also noticed that F1s of the trained mothers were larger in body size than the F1s of the naive mothers (Figure 2J). This intergenerational effect is consistent with previous findings showing that animals infected with PA14 retain their eggs more than worms raised on OP50 (Tan, Mahajan-Miklos, and Ausubel 1999) and thus their progeny are kept inside the mother until a more advanced stage. Therefore, we examined if the difference in body size (that could be due to their difference in the developmental stage) could be related with the difference in the food choice. We found no difference when comparing the body size between the F1s from the trained mothers in the “P0 low learning index” group and the F1s from the trained mothers in the “P0 high learning index” group (Figure 2K), as well as no correlation between the Parental experience_induced learning indexes of the F1s and the increase in body size in the progeny of the trained mothers compared with the progeny of the naive mothers (Figure 2L). In addition, we found that 8-hour exposure to PA14, while inducing robust avoidance of PA14 in P0 mothers did not alter the innate resistance to PA14 in the F1s (Supplementary Figure 3B). Together, these results show that the body size and the innate immune resistance do not play a significant role in regulating the parental experience-induced difference in offspring olfactory preference.

Next, we asked whether the learning effect in the P0 mothers trained with 8-hour exposure to PA14 regulates F2s. We isolated F2 embryos from F1s and cultivated F2s under the standard conditions on *E. coli* OP50. Similarly, we quantified the Parental experience_induced learning index in the F2s of each experiment and found no correlation with the same index of their F1 mothers (Supplementary Figure 4). These results indicate that the parental experience with PA14 only regulates the olfactory response of their first-generation progeny, potentially leaving the flexibility for F2s to respond to further changes in the environment.

Furthermore, we found that mutating *rrf-3* in *pk1426* also disrupted the Parental experience_induced behavioral change in the 8-hour training experiments. Despite the wild-type learning ability in the *rrf-3* P0 mothers (Figure 3A and 3B), the parental experience induced learning of the *rrf-3* F1s is no longer correlated with the learning of their mothers (Figure 3C). Together, these results show that the *rrf-3*-mediated endo-siRNA pathways also regulate intergenerational effects generated by training the P0 mothers for 8 hours. In addition, we found that during the two-choice assay, while the *rrf-3* F1s had a locomotory speed and a rate of reversals comparable to those in wild-type F1s, their rate of omega bends was lower than that in wild type (Figure 3D - 3F). These results identify the regulation of omega bends as the potential downstream effector of the *rrf-3*-mediated pathway in regulating intergenerational effects.

**Figure 3.**
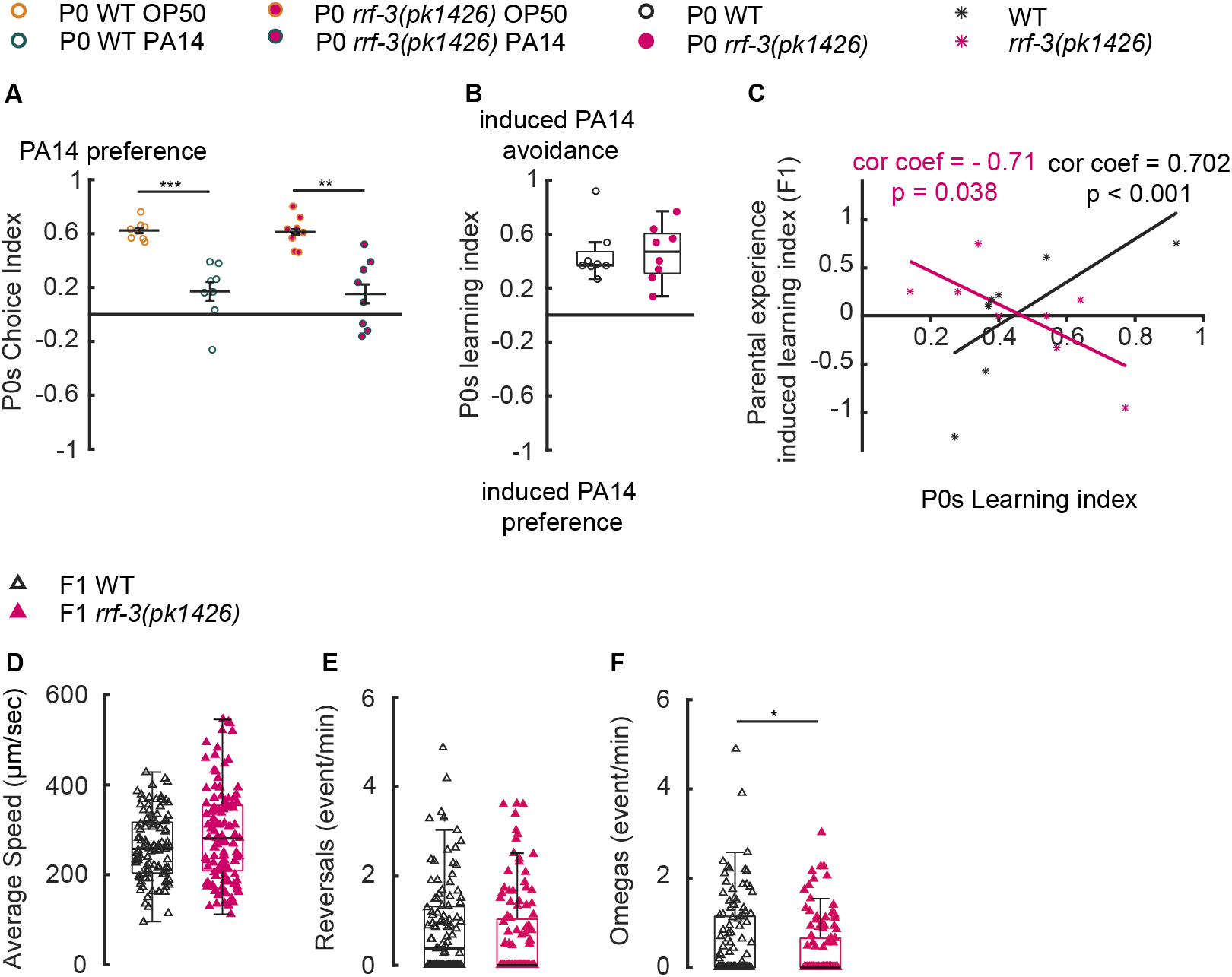
The RRF-3-pathway regulates intergenerational effects generated by 8-hour parental training. **(A, B, C)** The *rrf-3(pk1426)* mutant P0s display wild-type choice indexes **(A)** and learning **(B)** after 8-hour training with PA14; however, the parental experience-induced learning indexes in *rrf-3(pk1426)* mutant F1s do not positively correlate with their P0 mothers’ learning **(C)**. A two-tailed Student’s *t*-test with Bonferroni correction for multiple comparisons was used in **(A)** (n = 8 biological replicates each; *** p < 0.001, ** p < 0.01). A Wilcoxon rank sum test was used for comparisons in **(B)** (n = 8 biological replicates each, p > 0.05) and a permutation test was used to test linear correlation in (**C**). **(D - F)** The movement speed **(D)**, rate of reversals **(E)** and the rate of omega bends **(F)** in wild type (WT) F1s and *rrf-3(pk1426)* F1s. A Wilcoxon rank sum test was used for comparisons in (**D**) (p > 0.05), **(E)** (p > 0.05), and **(F)** (* p < 0.05). n = 107 worms for WT and n = 115 worms for *rrf-3(pk1426)*. Data is normally distributed in **A** and not normally distributed in **B - F**. In graph **A** horizontal bar represents Mean ± SEM and in box plots **(B, D - F)** dark horizontal bar represents median, box limits represent first and third quartile, and whiskers extend to values within 2.7 standard deviations. *** p < 0.001, ** p < 0.01, * p < 0.05. All data are included in the analysis.

## DISCUSSION

Previous studies on the effect of parental experience on the physiology and behavior of the offspring show that inter and transgenerational regulation is more complex than a unidirectional transfer of phenotypic traits acquired by the progenitors (Burton et al. 2017; Ng et al. 2010; Crews et al. 2014; Bohacek and Mansuy 2015; Kundakovic et al. 2013). Here, we find that a brief experience (4 hours) with the pathogenic strain of *Pseudomonas aeruginosa*, PA14, while reducing the preference for the odorants of the pathogen in the trained mothers, increases the preference in the progeny. Furthermore, extending the maternal exposure time to 8 hours switches the response of the progeny from attraction to avoidance. This effect is intergenerational and limited to only the progeny of the first generation. Recently it has been shown that an even longer exposure to the pathogenic bacteria (24 hours) not only induces avoidance in mothers but also in offspring for several generations (Moore et al. 2019). Together, these results propose that the same behavioral trait, pathogen avoidance acquired by the parents, can result from experiences of different levels of intensity that lead to diverse responses in progeny. It also suggests that the valence of the parental experience that is transmitted to the offspring is mostly mediated by the physiological, but not behavioral, response of the parents to the environmental conditions.

After 4-hour exposure to the pathogen, the *C. elegans* adults shift their preference towards food sources that are non-pathogenic (Zhang, Lu, and Bargmann 2005), a first line of behavioral defense towards increased infection. However, when presented with the option between PA14 and no food, worms still prefer PA14 (Y. Shen et al. 2014; Ha et al. 2010a). In addition, in the lawn-leaving assay where worms feed on a lawn of PA14 and gradually leave the lawn over time, most worms are still inside the lawn after consuming the pathogen for 4 hours and start to significantly leave the lawn after 8 hours (Singh and Aballay 2019). In such conditions, it is conceivable that after 4-hour exposure the food signal represented by PA14 is still significant in comparison with the signal of virulence and positively modulates the preference of the progeny towards the food source consumed by their mothers. Consistently, there are several reports showing that consumption of certain foods during gestation increases the preference of the progeny for the same foods after birth (Hepper et al. 2012; Crane et al. 2018; A. Liu and Urban 2017; Nehring et al. 2015). This modulation is likely to be adaptive since the food consumed by the mother is a potential source of food for the progeny. Extending the adult exposure to 8 hours enhances the innate immune response to PA14 (Troemel et al. 2006), which likely represents a more stably present adversity of virulence, resulting in the switch of offspring behavior from attraction to avoidance. An even longer exposure of 24 hours to a pathogenic lawn grown under conditions that enhance bacterial pathogenicity (Moore et al. 2019; LaBauve and Wargo, Matthew 2015) probably signals a constant presence of virulence that generates the transmission of the aversive information for several generations. Interestingly, the transgenerational modulation of metabolic traits in other species, including flies and rodents, is also influenced by the components of the parental diet, as well as the duration of consumption and the timing of exposure (Öst et al. 2014; Kundakovic et al. 2013; Palominos et al. 2017).

The *C. elegans rrf-3* is required for the generation of the 26G endo-siRNAs that are involved in the regulation of gene expression during spermatogenesis, oogenesis, as well as during zygotic development (Han et al. 2009). *rrf-3* mutants are depleted of the 26G RNA population (Han et al. 2009; Thivierge et al. 2012a), which affects endogenous gene expression during these processes. Thus, our results suggest that these *rrf-3*-mediated gene expression programs encode signals generated by parental learning to modulate offspring behavior. Furthermore, one of the *prg-2* transcripts encodes a homolog of a *C. elegans* Argonaute/Piwi-related protein PRG-1 that regulates piRNAs (Cox et al. 1998). Members of the piRNA pathway are expressed or enriched in the germline and are involved in maintaining germline genomic integrity by silencing non self-transcripts (Hashimshony et al. 2015; Weick and Miska 2014; Kasper, Gardner, and Reinke 2014). Despite the high level of similarity between *prg-1* and *prg-2*, the function of *prg-2* is not well understood (Das et al. 2008; Batista et al. 2008; Cox et al. 1998; Kasper, Gardner, and Reinke 2014). Our study reveals a role of *prg-2* in regulating parental learning induced modulation of olfactory preference. Thus, we show that both the endo-siRNA and piRNA pathways play a role in the modulation of olfactory preference in progeny after 4-hour exposure. Whether they affect the expression of the same genes is, however, an open question. Although the interaction between these pathways has been documented (Wang et al. 2014; E.-Z. Shen et al. 2018), the biogenesis of endo-siRNAs and piRNAs is different. At the level of behavioral regulation, while both mutations abolish the increased preference for PA14 in F1s as a result of training their mothers for 4 hours, their phenotypes are slightly different. This observation suggests that they may act in different ways when regulating parental experience-induced behavioral changes.

A major role of the endo-siRNA pathways is to modulate gene expression of endogenous genes (Han et al. 2009; Lee, Hammell, and Ambros 2006) – in its absence the basal level and the condition-dependent expression of these genes may be compromised. If parental experience induced changes in the progeny depend on the modulation of the gene expression in the progeny (Moore et al. 2019; Rechavi et al. 2014; Rodgers et al. 2013), the lack of the endo-siRNA pathways may compromise the maternal effects. Interestingly, a recent paper reports that the absence of the dsRNA-binding protein RDE-4 in neurons affects the pool of small RNAs in the germline in an heritable manner. Moreover, the expression of RDE-4 in the neurons of the great grandparents is critical for the normal behavior of the F3 progeny, revealing the role of RDE-4 in regulating the gene expression in the germline with an impact on the behavioral response of the future generations (Posner et al. 2019). RDE-4 acts upstream of both exo- and endo-siRNA pathways (Thivierge et al. 2012b; Vasale et al. 2010; Blanchard et al. 2011; Rechavi and Lev 2017). Several previous studies suggest that exogenous small RNAs can also regulate the physiology and behavior of the progeny by interacting with the exo-siRNA pathway in the germ line (Palominos et al. 2017; Kaletsky et al. 2019; De Abreu et al. 2019).

Together, the results from these and our study suggest that the normal function of the gene expression programs in the germline is important for parental experience-modulated sensory behaviors. It is intriguing that all of these different parental experiences depend on the small RNA pathways to generate modulatory effects on the offspring, revealing the flexibility of the small RNA-mediated gene expression programs to encode various environment conditions to regulate animal physiology across generations.

## ACKNOWLEDGMENTS

We thank *Caenorhabditis Genetics Center*, which is funded by National Institutes of Health - Office of Research Infrastructure Programs (P40 OD010440), for strains. Y.Z. is funded by National Institutes of Health.

## AUTHOR CONTRIBUITION

A.P., X.G., K.K. and Y.Z. designed the experiments, interpreted the results and wrote the manuscript. A.P., X.G. and K.K. performed the experiments and analyzed the data.

## Data Availability Statement

All the data generated and analyzed in this study are available upon request.

The method used for the automated assay has been previously published and the code is available upon request. The software used to analyze F1 chemotactic movement is available at https://www.mbfbioscience.com/wormlab and the Matlab code used for further analysis is available upon request.

## SUPPLEMENTARY MATERIAL

**Supplementary Figure 1.**
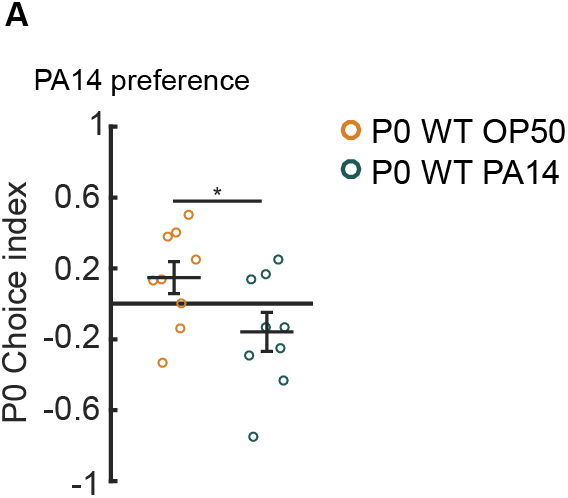
Transient aversive training with pathogenic bacterium PA14 for 4 hours decreases parental preference for PA14 odorants. **(A)** When tested in the two-choice plate assay, naive animals fed on *E. coli* OP50 (P0 WT OP50) prefer the odorants of PA14, while training with the pathogen (P0 WT PA14) for 4 hours decreases the preference. Thus, both the droplet assay and the two-choice plate assay similarly detect training-dependent change in PA14 preference. A two-tailed student’s *t*-test was used for comparison (n = 9 biological replicates, * p <0.05). Worms that freezed and stopped moving during the assay were considered to have a preference for OP50. Horizontal bars represent mean ± SEM and data are normally distributed. All the data are included in the analysis.

**Supplementary Figure 2.**
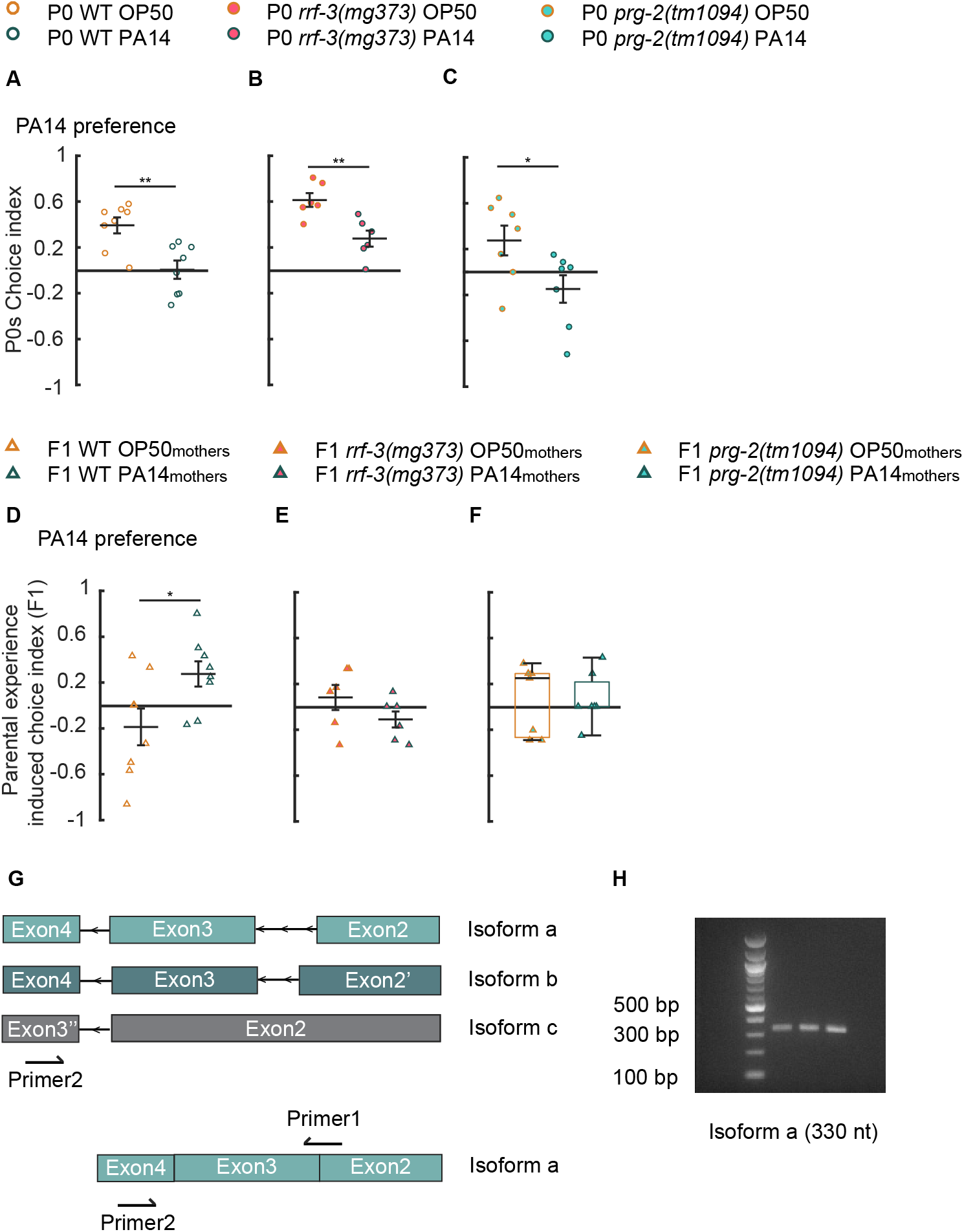
Mutants in the small RNA pathways affect offspring but not parental olfactory preference. **(A - F)** The missense mutation in *rrf-3(mg373)* **(B)** or the deletion in *prg-2(tm1094)* **(C)** does not abolish the decreased preference for PA14 in P0s after 4-hour training with the pathogen **(A-C);** however, they abolish the parental experience-induced PA14 preference in F1s (**D–F**). A two-tailed student’s *t*-test was used for comparisons in (**A**) (n = 8 biological replicates each, ** p < 0.01), (**B**) (n = 6 biological replicates each, ** p < 0.01), **(C)** (n = 7 biological replicates each, * p < 0.05), **(D)** (n = 8 biological replicates each, * p < 0.05) and **(E)** (n = 6 biological replicates each, p > 0.05). A Wilcoxon rank sum test was used for comparisons in **(F)** (n = 7 biological replicates each, p > 0.05). **(G, H)** Schematics of the genomic locus of *prg-2* showing 3 predicted isoforms of transcripts (1). Isoform a encodes a protein containing the PIWI and PAZ domains, while there is no predicted protein for isoform b or isoform c **(G)**. PCR products using primer 1 and primer 2, that are specific for isoform a of the *prg-2* gene, from a wild-type cDNA library (**H**). Three columns show the PRC products from increasing concentrations of the PCR template. Primer 1 (5’ CTCGTGCACTTCGAAAGGAA), Primer 2 (5’ CAGTCGGGAAGCACAATTCA), Data is normally distributed in **A - E** and not normally distributed in **F**. In graphs **A - E** horizontal bar represents Mean ± SEM and in box plots in **F** dark horizontal bar represents median, box limits represent first and third quartile, and whiskers extend to values within 2.7 standard deviations. ** p < 0.01, * p < 0.05. All data are included for analysis. (1) https://genome.ucsc.edu/cgi-bin/hgTracks?db=ce11&lastVirtModeType=default&lastVirtModeExtraState=&virtModeType=default&virtMode=0&nonVirtPosition=&position=chrIV%3A6526962%2D6529802&hgsid=783433341_2LhMwdEzSR6f8SgMApBbMQ5aX9Oa

**Supplementary Figure 3.**
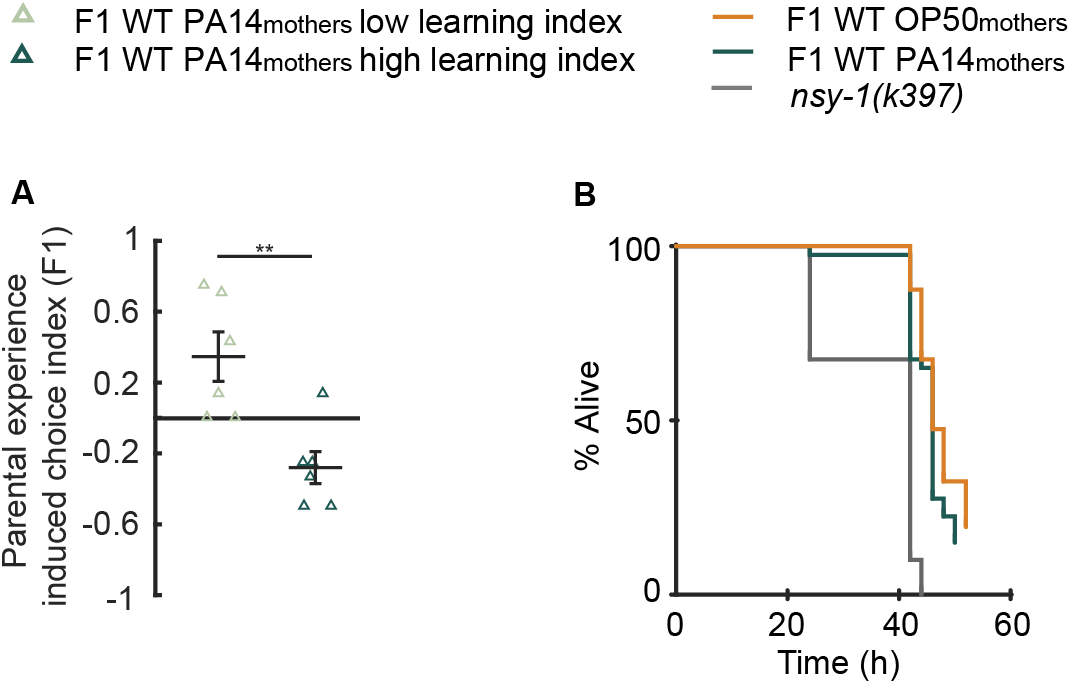
Extending parental training to 8 hours switches offspring response from attraction to aversion. **(A)** The parental experience-induced choice indexes in F1s whose mothers exhibit low learning indexes (light green triangles) are significantly more positive than the parental experience-induced choice indexes in F1s whose mothers exhibit high learning indexes (dark green triangles). A two-tailed student’s *t*-test was used for comparisons (n = 6 biological replicates for each, ** p < 0.01). Data is normally distributed and horizontal bars represents Mean ± SEM. **(B)** Parental learning experience does not alter innate immune resistance to PA14. *nsy-1(ky397)* mutant animals that display reduced resistance are used as experimental controls [S1]. Mantel-Cox log rank test was used to test for survival rate (n = 2 biological replicates, F1 WT OP50_mothers_ *vs* F1 WT PA14_mothers_, p = 0.104, F1 WT OP50_mothers_ *vs nsy-1(ky397)*, p < 0.001).

**Supplementary Figure 4.**
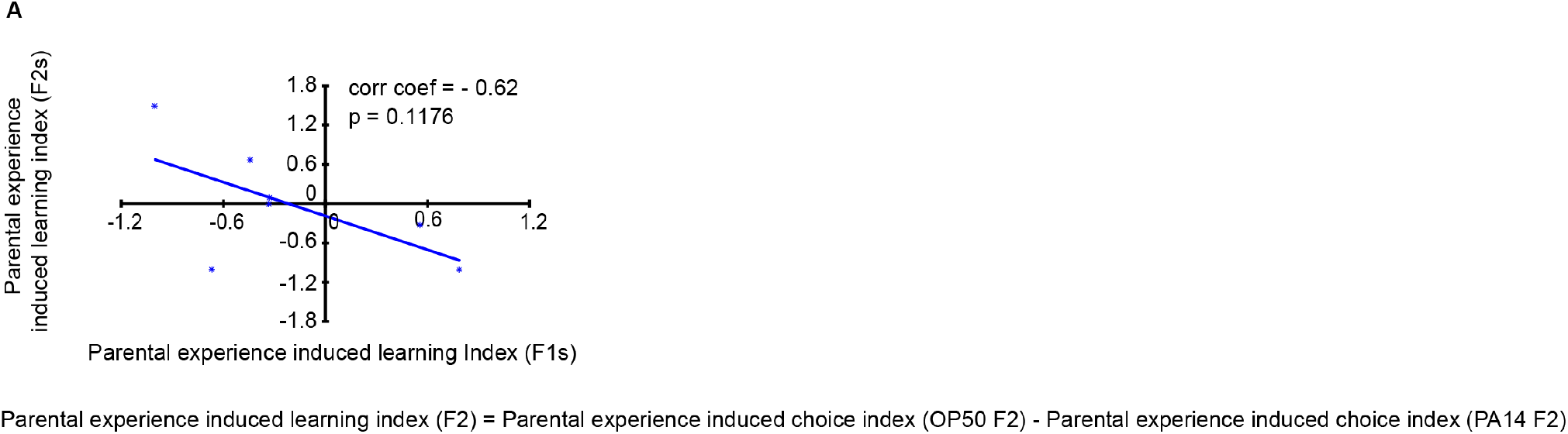
The modulation of olfactory choice in the F1 progeny is not transmitted to the second generation (F2s). **(A)** Parental experience-induced learning indexes in F2s are not significantly correlated with the parental experience-induced learning indexes in the F1s. Permutation test was used to test for linear correlation (n = 5 biological replicates).

## REFERENCES

Abreu, Diana Andrea Fernandes De, Thalia Salinas-Giegé, Laurence Maréchal-Drouard, and Jean-Jacques Remy. 2019. “Alanine TRNA Translate Environment into Behavior in *Caenorhabditis Elegans*.” SSRN Electronic Journal. https://doi.org/10.2139/ssrn.3310818.

Agrawal, Anurag A., Christian Laforsch, and Ralph Tollrian. 1999. “Transgenerational Induction of Defences in Animals and Plants.” Nature 401 (6748): 60–63. https://doi.org/10.1038/43425.

Ashe, Alyson, Alexandra Sapetschnig, Eva Maria Weick, Jacinth Mitchell, Marloes P. Bagijn, Amy C. Cording, Anna Lisa Doebley, et al. 2012. “PiRNAs Can Trigger a Multigenerational Epigenetic Memory in the Germline of C. Elegans.” Cell 150 (1): 88–99. https://doi.org/10.1016/j.cell.2012.06.018.

Bagijn, Marloes P., Leonard D. Goldstein, Alexandra Sapetschnig, Eva Maria Weick, Samir Bouasker, Nicolas J. Lehrbach, Martin J. Simard, and Eric A. Miska. 2012. “Function, Targets, and Evolution of Caenorhabditis Elegans PiRNAs.” Science. https://doi.org/10.1126/science.1220952.

Batista, Pedro J., J. Graham Ruby, Julie M. Claycomb, Rosaria Chiang, Noah Fahlgren, Kristin D. Kasschau, Daniel A. Chaves, et al. 2008. “PRG-1 and 21U-RNAs Interact to Form the PiRNA Complex Required for Fertility in C. Elegans.” Molecular Cell 31 (1): 67–78. https://doi.org/10.1016/j.molcel.2008.06.002.

Belicard, Tony, Pree Jareosettasin, and Peter Sarkies. 2018. “The PiRNA Pathway Responds to Environmental Signals to Establish Intergenerational Adaptation to Stress.” BMC Biology 16 (1): 1–14. https://doi.org/10.1186/s12915-018-0571-y.

Blanchard, Daniel, Poornima Parameswaran, Javier Lopez-Molina, Jonathan Gent, Jamie Fleenor Saynuk, and Andrew Fire. 2011. “On the Nature of in Vivo Requirements for Rde-4 in RNAi and Developmental Pathways in C. Elegans.” RNA Biology 8 (3): 458–67. https://doi.org/10.4161/rna.8.3.14657.

Bohacek, Johannes, and Isabelle M. Mansuy. 2015. “Molecular Insights into Transgenerational Non-Genetic Inheritance of Acquired Behaviours.” Nature Reviews Genetics 16 (11): 641–52. https://doi.org/10.1038/nrg3964.

Bond, Donna M., and David C. Baulcombe. 2014. “Small RNAs and Heritable Epigenetic Variation in Plants.” Trends in Cell Biology 24 (2): 100–107. https://doi.org/10.1016/j.tcb.2013.08.001.

Bozler, Julianna, Balint Z. Kacsoh, and Giovanni Bosco. 2019. “Transgenerational Inheritance of Ethanol Preference Is Caused by Maternal NPF Repression.” ELife 8: 1–18. https://doi.org/10.7554/eLife.45391.

Brennecke, Julius. 2008. “An Epigenetic Role for Maternally Inherited PiRNAs in Transposon Silencing.” Science 1387 (November): 1387–92. https://doi.org/10.1126/science.1165171.

Brenner, S. 1974. “The Genetics of Caenorhabditis Elegans.” Genetics.

Burton, Nicholas O., Tokiko Furuta, Amy K. Webster, Rebecca E.W. Kaplan, L. Ryan Baugh, Swathi Arur, and H. Robert Horvitz. 2017. “Insulin-like Signalling to the Maternal Germline Controls Progeny Response to Osmotic Stress.” Nature Cell Biology 19 (3): 252–57. https://doi.org/10.1038/ncb3470.

Claycomb, Julie M., Pedro J. Batista, Ka Ming Pang, Weifeng Gu, Jessica J. Vasale, Josien C. van Wolfswinkel, Daniel A. Chaves, et al. 2009. “The Argonaute CSR-1 and Its 22G-RNA Cofactors Are Required for Holocentric Chromosome Segregation.” Cell 139 (1): 123–34. https://doi.org/10.1016/j.cell.2009.09.014.

Cox, Daniel N., Anna Chao, Jeff Baker, Lisa Chang, Dan Qiao, and Haifan Lin. 1998. “A Novel Class of Evolutionarily Conserved Genes Defined by Piwi Are Essential for Stem Cell Self-Renewal.” Genes and Development 12 (23): 3715–27. https://doi.org/10.1101/gad.12.23.3715.

Crane, Adam L., Emilee J. Helton, Maud C.O. Ferrari, and Alicia Mathis. 2018. “Learning to Find Food: Evidence for Embryonic Sensitization and Juvenile Social Learning in a Salamander.” Animal Behaviour 142: 199–206. https://doi.org/10.1016/j.anbehav.2018.06.021.

Crews, David, Ross Gillette, Isaac Miller-Crews, Andrea C. Gore, and Michael K. Skinner. 2014. “Nature, Nurture and Epigenetics.” Molecular and Cellular Endocrinology 398 (1–2): 42–52. https://doi.org/10.1016/j.mce.2014.07.013.

Das, Partha P., Marloes P. Bagijn, Leonard D. Goldstein, Julie R. Woolford, Nicolas J. Lehrbach, Alexandra Sapetschnig, Heeran R. Buhecha, et al. 2008. “Piwi and PiRNAs Act Upstream of an Endogenous SiRNA Pathway to Suppress Tc3 Transposon Mobility in the Caenorhabditis Elegans Germline.” Molecular Cell 31 (1): 79–90. https://doi.org/10.1016/j.molcel.2008.06.003.

Deas, Joseph B., Leo Blondel, and Cassandra G. Extavour. 2019. “Ancestral and Offspring Nutrition Interact to Affect Life-History Traits in Drosophila Melanogaster.” Proceedings of the Royal Society B: Biological Sciences 286 (1897). https://doi.org/10.1098/rspb.2018.2778.

Dew-Budd, Kelly, Julie Jarnigan, and Laura K. Reed. 2016. “Genetic and Sex-Specific Transgenerational Effects of a High Fat Diet in Drosophila Melanogaster.” PLoS ONE 11 (8): 1–20. https://doi.org/10.1371/journal.pone.0160857.

Dias, Brian G., and Kerry J. Ressler. 2014. “Parental Olfactory Experience Influences Behavior and Neural Structure in Subsequent Generations.” Nature Neuroscience 17 (1): 89–96. https://doi.org/10.1038/nn.3594.

Gapp, Katharina, Ali Jawaid, Peter Sarkies, Johannes Bohacek, Pawel Pelczar, Julien Prados, Laurent Farinelli, Eric Miska, and Isabelle M. Mansuy. 2014. “Implication of Sperm RNAs in Transgenerational Inheritance of the Effects of Early Trauma in Mice.” Nature Neuroscience 17 (5): 667–69. https://doi.org/10.1038/nn.3695.

Garcia, John, Walter G. Hankins, and Kenneth W. Rusiniak. 1974. “Behavioral Regulation of the Milieu Interne in Man and Rat.” Science. https://doi.org/10.1126/science.185.4154.824.

Gent, Jonathan I., Mara Schvarzstein, Anne M. Villeneuve, Sam Guoping Gu, Verena Jantsch, Andrew Z. Fire, and Antoine Baudrimont. 2009. “A Caenorhabditis Elegans RNA-Directed RNA Polymerase in Sperm Development and Endogenous RNA Interference.” Genetics 183 (4): 1297–1314. https://doi.org/10.1534/genetics.109.109686.

Gent, Jonathan I, Ayelet T Lamm, Derek M Pavelec, Jay M Maniar, Poornima Parameswaran, Li Tao, Scott Kennedy, and Andrew Z Fire. 2010. “Distinct Phases of SiRNA Synthesis in an Endogenous RNAi Pathway in C. Elegans Soma.” Molecular Cell 37 (5): 679–89. https://doi.org/10.1016/j.molcel.2010.01.012.

Ha, Heon ick, Michael Hendricks, Yu Shen, Christopher V. Gabel, Christopher Fang-Yen, Yuqi Qin, Daniel Colón-Ramos, Kang Shen, Aravinthan D.T. Samuel, and Yun Zhang. 2010a. “Functional Organization of a Neural Network for Aversive Olfactory Learning in Caenorhabditis Elegans.” Neuron 68 (6): 1173–86. https://doi.org/10.1016/j.neuron.2010.11.025.

Han, Ting, Arun Prasad, Tim T Harkins, Pascal Bouffard, Colin Fitzpatrick, and Diana S Chu. 2009. “26G Endo-SiRNAs Regulate Spermatogenic and Zygotic Gene Expression in Caenorhabditis Elegans,” 1–6.

Hashimshony, Tamar, Martin Feder, Michal Levin, Brian K. Hall, and Itai Yanai. 2015. “Spatiotemporal Transcriptomics Reveals the Evolutionary History of the Endoderm Germ Layer.” Nature. https://doi.org/10.1038/nature13996.

Hepper, P. G., D. L. Wells, S. Millsopp, K. Kraehenbuehl, S. A. Lyn, and O. Mauroux. 2012. “Prenatal and Early Sucking Influences on Dietary Preference in Newborn, Weaning, and Young Adult Cats.” Chemical Senses 37 (8): 755–66. https://doi.org/10.1093/chemse/bjs062.

Kaletsky, Rachel, Rebecca S Moore, Lance L Parsons, and Coleen T Murphy. 2019. “Cross-Kingdom Recognition of Bacterial Small RNAs Induces Transgenerational Pathogenic Avoidance.”

Kasper, Dionna M., Kathryn E. Gardner, and Valerie Reinke. 2014. “Homeland Security in the C. Elegans Germ Line: Insights into the Biogenesis and Function of Pirnas.” Epigenetics 9 (1): 62–74. https://doi.org/10.4161/epi.26647.

Kim, Dennis H., Rhonda Feinbaum, Geneviève Alloing, Fred E. Emerson, Danielle A. Garsin, Hideki Inoue, Miho Tanaka-Hino, et al. 2002. “A Conserved P38 MAP Kinase Pathway in Caenorhabditis Elegans Innate Immunity.” Science. https://doi.org/10.1126/science.1073759.

Kundakovic, Marija, Kathryn Gudsnuk, Becca Franks, Jesus Madrid, Rachel L. Miller, Frederica P. Perera, and Frances A. Champagne. 2013. “Sex-Specific Epigenetic Disruption and Behavioral Changes Following Low-Dose in Utero Bisphenol a Exposure.” Proceedings of the National Academy of Sciences of the United States of America 110 (24): 9956–61. https://doi.org/10.1073/pnas.1214056110.

LaBauve, Annette. E., and J. Wargo, Matthew. 2015. “Growth and Laboratory Maintenance of Pseudomonas Aeruginosa.” Current Protocol Microbiology May: 1–11. https://doi.org/10.1002/9780471729259.mc06e01s25.Growth.

Lee, Rosalind C., Christopher M. Hammell, and Victor Ambros. 2006. “Interacting Endogenous and Exogenous RNAi Pathways in Caenorhabditis Elegans.” Rna 12 (4): 589–97. https://doi.org/10.1261/rna.2231506.

Liu, Annie, and Nathaniel N. Urban. 2017. “Prenatal and Early Postnatal Odorant Exposure Heightens Odor-Evoked Mitral Cell Responses in the Mouse Olfactory Bulb.” Eneuro 4 (5): ENEURO.0129-17.2017. https://doi.org/10.1523/ENEURO.0129-17.2017.

Liu, He, Wenxing Yang, Taihong Wu, Fengyun Duan, Edward Soucy, Xin Jin, and Yun Zhang. 2018. “Cholinergic Sensorimotor Integration Regulates Olfactory Steering.” Neuron 97 (2): 390–405.e3. https://doi.org/10.1016/j.neuron.2017.12.003.

Moore, Rebecca S, Rachel Kaletsky, Coleen T Murphy, Rebecca S Moore, Rachel Kaletsky, and Coleen T Murphy. 2019. “Piwi / PRG-1 Argonaute and TGF-b Mediate Transgenerational Learned Pathogenic Avoidance.” Cell 177 (7): 1827–1841.e12. https://doi.org/10.1016/j.cell.2019.05.024.

Mueller, Bridget R., and Tracy L. Bale. 2006. “Impact of Prenatal Stress on Long Term Body Weight Is Dependent on Timing and Maternal Sensitivity.” Physiology and Behavior 88 (4–5): 605–14. https://doi.org/10.1016/j.physbeh.2006.05.019.

Nehring, Ina, Tanja Kostka, Rüdiger von Kries, and Eva A Rehfuess. 2015. “Impacts of In Utero and Early Infant Taste Experiences on Later Taste Acceptance: A Systematic Review.” The Journal of Nutrition 145 (6): 1271–79. https://doi.org/10.3945/jn.114.203976.

Ng, Sheau Fang, Ruby C.Y. Lin, D. Ross Laybutt, Romain Barres, Julie A. Owens, and Margaret J. Morris. 2010. “Chronic High-Fat Diet in Fathers Programs β 2-Cell Dysfunction in Female Rat Offspring.” Nature 467 (7318): 963–66. https://doi.org/10.1038/nature09491.

Okamura, Katsutomo, and Eric C. Lai. 2008. “Endogenous Small Interfering RNAs in Animals.” Nature Reviews Molecular Cell Biology 9 (9): 673–78. https://doi.org/10.1038/nrm2479.

Öst, Anita, Adelheid Lempradl, Eduard Casas, Melanie Weigert, Theodor Tiko, Merdin Deniz, Lorena Pantano, et al. 2014. “Paternal Diet Defines Offspring Chromatin State and Intergenerational Obesity.” Cell 159 (6): 1352–64. https://doi.org/10.1016/j.cell.2014.11.005.

Painter, Rebecca C., Tessa J. Roseboom, and Otto P. Bleker. 2005. “Prenatal Exposure to the Dutch Famine and Disease in Later Life: An Overview.” Reproductive Toxicology 20 (3): 345–52. https://doi.org/10.1016/j.reprotox.2005.04.005.

Palominos, M. Fernanda, Lidia Verdugo, Carolaing Gabaldon, Bernardo Pollak, Javiera Ortíz-Severín, Macarena A. Varas, Francisco P. Chávez, and Andrea Calixto. 2017. “Transgenerational Diapause as an Avoidance Strategy against Bacterial Pathogens in Caenorhabditis Elegans.” MBio 8 (5): 1–18. https://doi.org/10.1128/mBio.01234-17.

Pavelec, Derek M., Jennifer Lachowiec, Thomas F. Duchaine, Harold E. Smith, and Scott Kennedy. 2009. “Requirement for the ERI/DICER Complex in Endogenous RNA Interference and Sperm Development in Caenorhabditis Elegans.” Genetics 183 (4): 1283–95. https://doi.org/10.1534/genetics.109.108134.

Pierce-Shimomura, Jonathan T., Thomas M. Morse, and Shawn R. Lockery. 1999. “The Fundamental Role of Pirouettes in Caenorhabditis Elegans Chemotaxis.” Journal of Neuroscience. https://doi.org/10.1523/jneurosci.19-21-09557.1999.

Posner, Rachel, Itai Antoine Toker, Olga Antonova, Ekaterina Star, Sarit Anava, Eran Azmon, Michael Hendricks, Shahar Bracha, Hila Gingold, and Oded Rechavi. 2019. “Neuronal Small RNAs Control Behavior Transgenerationally.” Cell 177 (7): 1814–1826.e15. https://doi.org/10.1016/j.cell.2019.04.029.

Rankin, Catharine H. 2015. “A Review of Transgenerational Epigenetics for RNAi, Longevity, Germline Maintenance and Olfactory Imprinting in Caenorhabditis Elegans.” Journal of Experimental Biology 218 (1): 41–49. https://doi.org/10.1242/jeb.108340.

Rechavi, Oded, Leah Houri-Ze’Evi, Sarit Anava, Wee Siong Sho Goh, Sze Yen Kerk, Gregory J. Hannon, and Oliver Hobert. 2014. “Starvation-Induced Transgenerational Inheritance of Small RNAs in C. Elegans.” Cell 158 (2): 277–87. https://doi.org/10.1016/j.cell.2014.06.020.

Rechavi, Oded, and Itamar Lev. 2017. “Principles of Transgenerational Small RNA Inheritance in Caenorhabditis Elegans.” Current Biology 27 (14): R720–30. https://doi.org/10.1016/j.cub.2017.05.043.

Remy, Jean-jacques. 2010. “Stable Inheritance of an Acquired Behavior in Caenorhabditis Elegans.” Current Biology 20 (20): 3–5. https://doi.org/10.1016/j.cub.2010.08.013.

Rodgers, Ali B., Christopher P. Morgan, Stefanie L. Bronson, Sonia Revello, and Tracy L. Bale. 2013. “Paternal Stress Exposure Alters Sperm MicroRNA Content and Reprograms Offspring HPA Stress Axis Regulation.” Journal of Neuroscience 33 (21): 9003–12. https://doi.org/10.1523/JNEUROSCI.0914-13.2013.

Samuel, Buck S., Holli Rowedder, Christian Braendle, Marie Anne Félix, and Gary Ruvkun. 2016. “Caenorhabditis Elegans Responses to Bacteria from Its Natural Habitats.” Proceedings of the National Academy of Sciences of the United States of America 113 (27): E3941–49. https://doi.org/10.1073/pnas.1607183113.

Shen, En-Zhi, H Chen, Ahmet R Ozturk, Shikui Tu, M Shirayama, W Tang, Yue-He Ding, Si-Yuan Dai, Zhiping Weng, and Craig C Mello. 2018. “Identification of PiRNA Binding Sites Reveals the Argonaute Regulatory Landscape of the C Elegans Germline.” Cell 172 (3): 937–51. https://doi.org/10.1016/j.physbeh.2017.03.040.

Shen, Y., J. Zhang, J. A. Calarco, and Y. Zhang. 2014. “EOL-1, the Homolog of the Mammalian Dom3Z, Regulates Olfactory Learning in C. Elegans.” Journal of Neuroscience 34 (40): 13364–70. https://doi.org/10.1523/JNEUROSCI.0230-14.2014.

Simmer, Femke, Marcel Tijsterman, Susan Parrish, Sandhya P. Koushika, Michael L. Nonet, Andrew Fire, Julie Ahringer, and Ronald H.A. Plasterk. 2002. “Loss of the Putative RNA-Directed RNA Polymerase RRF-3 Makes C. Elegans Hypersensitive to RNAi.” Current Biology 12 (15): 1317–19. https://doi.org/10.1016/S0960-9822(02)01041-2.

Singh, Jogender, and Alejandro Aballay. 2019. “Microbial Colonization Activates an Immune Fight- and-Flight Response via Neuroendocrine Signaling.” Developmental Cell 49 (1): 89–99.e4. https://doi.org/10.1016/j.devcel.2019.02.001.

Tan, M W, S Mahajan-Miklos, and F M Ausubel. 1999. “Killing of *Caenorhabditis Elegans* by *Pseudomonas Aeruginosa* Used to Model Mammalian Bacterial Pathogenesis.” Proceedings of the National Academy of Sciences of the United States of America 96 (2): 715–20. https://doi.org/10.1073/pnas.96.2.715.

Thivierge, Caroline, Neetha Makil, Mathieu Flamand, Jessica J. Vasale, Craig C. Mello, James Wohlschlegel, Darryl Conte, and Thomas F. Duchaine. 2012a. “Tudor Domain ERI-5 Tethers an RNA-Dependent RNA Polymerase to DCR-1 to Potentiate Endo-RNAi.” Nature Structural and Molecular Biology 19 (1): 90–98. https://doi.org/10.1038/nsmb.2186.

Troemel, Emily R., Stephanie W. Chu, Valerie Reinke, Siu Sylvia Lee, Frederick M. Ausubel, and Dennis H. Kim. 2006. “P38 MAPK Regulates Expression of Immune Response Genes and Contributes to Longevity in C. Elegans.” PLoS Genetics. https://doi.org/10.1371/journal.pgen.0020183.

Vasale, Jessica J., Weifeng Gu, Caroline Thivierge, Pedro J. Batista, Julie M. Claycomb, Elaine M. Youngman, Thomas F. Duchaine, Craig C. Mello, and Darryl Conte. 2010. “Sequential Rounds of RNA-Dependent RNA Transcription Drive Endogenous Small-RNA Biogenesis in the ERGO-1/ Argonaute Pathway.” Proceedings of the National Academy of Sciences of the United States of America 107 (8): 3582–87. https://doi.org/10.1073/pnas.0911908107.

Wang, Jia Jia, Dong Ya Cui, Tengfei Xiao, Xubin Sun, Peng Zhang, Runsheng Chen, Shunmin He, and Da Wei Huang. 2014. “The Influences of PRG-1 on the Expression of Small RNAs and MRNAs.” BMC Genomics 15 (1): 1–11. https://doi.org/10.1186/1471-2164-15-321.

Weick, Eva Maria, and Eric A. Miska. 2014. “PiRNAs: From Biogenesis to Function.” Development (Cambridge) 141 (18): 3458–71. https://doi.org/10.1242/dev.094037.

Wells, Jonathan C K. 2007. “The Thrifty Phenotype as an Adaptive Maternal Effect” 82: 143–72. https://doi.org/10.1111/j.1469-185X.2006.00007.x.

Youngman, Elaine M., and Julie M. Claycomb. 2014. “From Early Lessons to New Frontiers: The Worm as a Treasure Trove of Small RNA Biology.” Frontiers in Genetics 5 (NOV): 1–13. https://doi.org/10.3389/fgene.2014.00416.

Zhang, Yun, Hang Lu, and Cornelia I. Bargmann. 2005. “Pathogenic Bacteria Induce Aversive Olfactory Learning in Caenorhabditis Elegans.” Nature 438 (7065): 179–84. https://doi.org/10.1038/nature04216.

## Supplementary References

S1. Kim, D.H., Feinbaum, R., Alloing, G., Emerson, F.E., Garsin, D.A., Inoue, H., Tanaka-Hino, M., Hisamoto, N., Matsumoto, K., Tan, M.W., et al. (2002). A conserved p38 MAP kinase pathway in Caenorhabditis elegans innate immunity. Science 297, 623–6.

